# Gonadal histology and variation in gonadosomatic index in captive moonfish *Mene maculata*

**DOI:** 10.64898/2026.07.12.738070

**Authors:** Masato Oi, Chiho Ogawa, Kenichi Fujii, Toshiaki Mori, Sayaka Matsuo, Kazuya Fukuda

## Abstract

The moonfish *Mene maculata* (Bloch and Schneider, 1801), the sole extant species of the family Menidae, is widely distributed in the Indo-West Pacific. Although its spawning season has previously been inferred from seasonal variation in the gonadosomatic index (GSI), the histological basis for interpreting GSI as an indicator of spawning-capable status remains limited. Here we describe the gonadal structure of male and female *M. maculata* and evaluate how GSI and standard length relate to spawning-capable status based on germ cell development. The testis was lobular and exhibited an unrestricted spermatogonial distribution, a structure widely observed among neoteleosts, indicating that the phylogenetic distinctiveness of *M. maculata* was not associated with a distinctive testicular structure. In the spawning-capable female, oocytes at multiple developmental stages, from primary growth to oocyte maturation, co-occurred within the ovary, indicating asynchronous ovarian development. This finding suggests that *M. maculata* may be a batch spawner rather than a total spawner as previously inferred. The spawning-capable female had a GSI consistent with previously inferred spawning season estimates. By contrast, histologically examined males were spawning capable at GSI values lower than those previously associated with the inferred spawning season, suggesting that male spawning-capable status may persist beyond the period inferred from elevated GSI alone. This study provides preliminary histological information on gonadal development in captive *M. maculata* and context for interpreting previously published field measurements of GSI.

## Introduction

Fishes are among the most species-rich groups of vertebrates, with more than 37,000 valid species currently recognized (Fricke et al. 2026). This diversity includes many phylogenetically distinctive, species-poor lineages, including monotypic families represented by a single extant species. Because such lineages often possess unique morphological, ecological, and life-history traits, they provide important opportunities for understanding the diversity and evolutionary processes of fishes. One example is the naked sand lance *Hypoptychus dybowskii* Steindachner, 1880, a gasterosteiform fish from a species-poor lineage whose reproductive ecology has been described. This species shows an unusual reproductive behavior among gasterosteiform fishes: females deposit egg masses in coils around the branching points of sargasso weeds, and males pick at the egg masses with the snout after spawning, thereby attaching them firmly around the weeds, but do not perform further paternal care such as fanning (Akagawa and Okiyama 1993). Characterizing the biological traits of such distinctive lineages is therefore essential for understanding fish diversity.

Gonadal structure, a key reproductive trait, has been studied comparatively not only in relation to reproductive ecology but also as a character relevant to teleost phylogeny and evolution (Grier 1981; Parenti and Grier 2004; Cole 2010; Uribe et al. 2014). In teleosts, testes are broadly classified into tubular and lobular types on the basis of their structural organization. Lobular testes are further classified into unrestricted and restricted spermatogonial types according to the distribution of spermatogonia, and these types occur unevenly among teleost lineages (Grier 1981; Uribe et al. 2014). Similarly, ovarian structure and the synchrony of oocyte development are closely linked to spawning pattern and fecundity type (Wallace and Selman 1981; Murua and Saborido-Rey 2003). In species with synchronous ovarian development, oocytes mature as a single cohort and are typically released in one spawning event; these species are termed total spawners. By contrast, batch spawners, which spawn repeatedly within a spawning season, may retain oocytes at several developmental stages simultaneously within the ovary. In particular, asynchronous ovarian development, in which oocytes from the primary growth phase through multiple vitellogenic stages co-occur, is generally associated with batch spawning characterized by continuous oocyte recruitment and repeated spawning throughout the spawning season (Wallace and Selman 1981; Murua and Saborido-Rey 2003; Brown-Peterson et al. 2011). Clarifying gonadal structure and gametogenesis can therefore provide insights into both reproductive ecology and evolutionary history, and is especially valuable for phylogenetically distinctive lineages for which reproductive records remain limited.

The moonfish *Mene maculata* (Bloch and Schneider, 1801) is the sole extant species of the family Menidae. It occurs at depths of 50–200 m and is widely distributed in the Indo-West Pacific, from East Africa to southern Japan, northeastern Australia, and New Caledonia (Senou 2013), and is an important fishery target in several countries. Its spawning season has been investigated in specimens collected from the northern South China Sea (Du et al. 2012) and off the southern coast of Java (Gani et al. 2024). Based on monthly changes in the gonadosomatic index (GSI) and macroscopic assessment of gonad maturation, the spawning peak has been estimated to occur around August–October. From oocyte diameter composition, Gani et al. (2024) further inferred that the species is a total spawner. However, information on the reproductive biology of *M. maculata* remains limited, and the relationship between GSI, germ cell development, and spawning-capable status has not been examined histologically. Histological assessment of germ cell development in relation to GSI would help identify the GSI range associated with the spawning-capable phase and provide a more robust basis for interpreting the reproductive ecology of this species.

In this study, we histologically describe the testicular and ovarian structure of *M. maculata* and assess spawning-capable status on the basis of germ cell developmental stage. We further examine how GSI and standard length are associated with histologically determined spawning-capable status in each sex. Based on these results, we provide a histological basis for interpreting previous spawning season estimates derived from seasonal changes in GSI and macroscopic gonad maturity.

## Materials and methods

### Fish collection and sampling

All fish used in this study were collected using a set net and light traps in Kuji Bay, Amami-Oshima, Kagoshima Prefecture, Japan, in July 2021. To minimize transport stress, the fish were held for 28 days in an enclosure net at the capture site and fed krill (Fig. 1a). They were then transported to Aquamarine Fukushima, Fukushima Prefecture, Japan, and maintained in three tanks (43, 24, and 59 t) at appropriate stocking densities (Fig. 1b). Water temperature was maintained at 19.2–28.8 °C under a 10 h light:14 h dark photoperiod. Fish were fed silver-stripe round herring *Spratelloides gracilis*, krill, and Sakura shrimp *Lucensosergia lucens* three to five times daily.

**Fig. 1.**
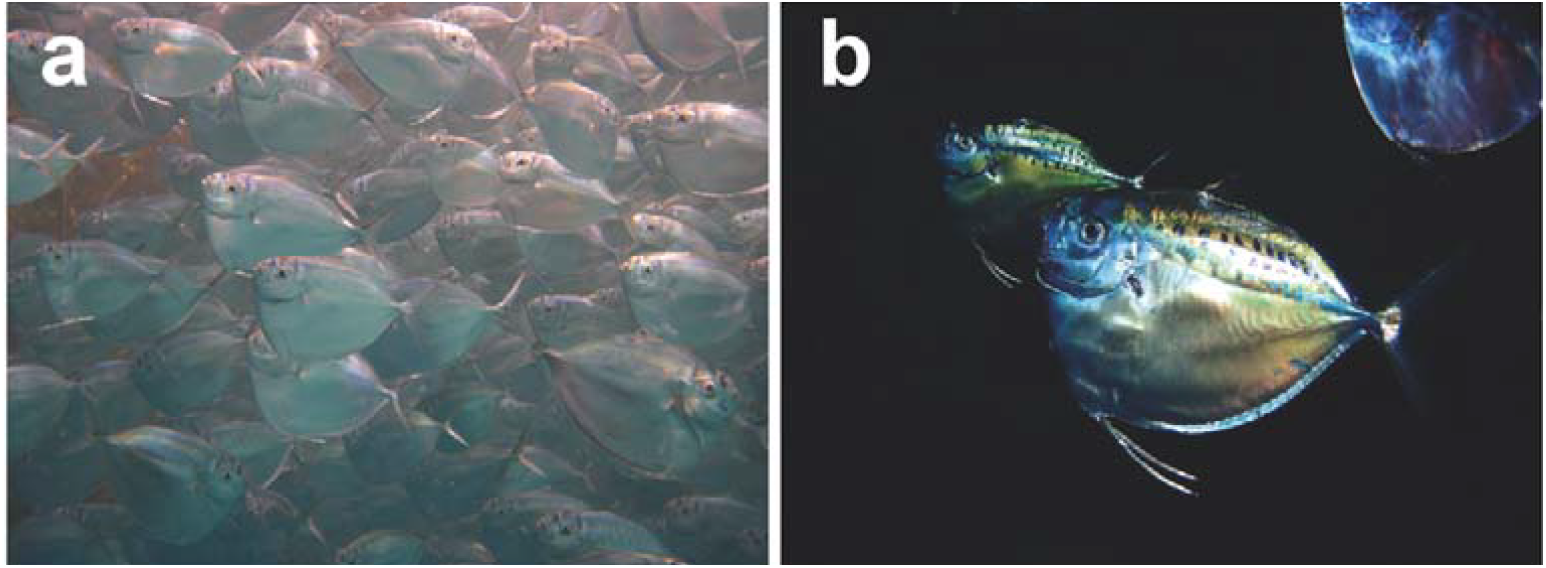
Photographs of live Mene maculata used in this study. (**a**) Individuals held in an enclosure net at the capture site after collection. (**b**) Individuals maintained in a rearing tank at Aquamarine Fukushima, Fukushima Prefecture, Japan, after transport from the capture site

Data from 98 of the 133 fish involved in transport and aquarium rearing between 2021 and 2023 were used in this study. These 98 fish comprised 95 that died either during transport (*n* = 10) or during tank rearing (*n* = 85), and three that were euthanized for sampling. During tank rearing, fish were checked daily as part of routine aquarium husbandry. Although the exact times of death were unknown, the interval between death and detection was estimated not to exceed 24 h based on these daily checks. When dead fish were found, their standard length (SL) and whole-body weight (BW) were measured before dissection. At dissection, the gonads retained their macroscopic morphology. Gonad weight (GW) was then measured. Gonads from nine fish that died during rearing between February and October 2023 were fixed in 10% formalin in seawater (FUJIFILM Wako Pure Chemical Corporation, Osaka, Japan) for histological analysis. To obtain histological material in a fresher condition, three additional live fish were sampled in April 2023. These fish were deeply anesthetized and euthanized with an overdose of 2-phenoxyethanol (FUJIFILM Wako Pure Chemical Corporation, Osaka, Japan). SL, BW, and GW were then measured, and the gonads were fixed in Bouin’s solution.

### Gonadal histology

Fixed gonads were embedded in paraffin (Leica Biosystems, Nussloch, Germany), sectioned at 5 µm, and stained with Mayer’s hematoxylin and eosin Y (FUJIFILM Wako Pure Chemical Corporation, Osaka, Japan). Histological images were obtained using a BX53 microscope and a DP74 camera (Evident, Tokyo, Japan). Developmental stages of germ cells and associated terminology were identified with reference to Grier (1981), Wallace and Selman (1981), Tyler and Sumpter (1996), Murua and Saborido-Rey (2003), Schulz et al. (2010), Brown-Peterson et al. (2011), Uribe et al. (2014), Saber et al. (2015), and Fish et al. (2020). Assessment of spawning-capable status was based primarily on the criteria of Brown-Peterson et al. (2011). In males, individuals with spermatozoa present in the lumen of the lobules and in the sperm ducts were considered to be in the spawning-capable phase. In females, individuals with tertiary vitellogenic oocytes or oocytes undergoing early stages of oocyte maturation, such as germinal vesicle migration, were considered to be in the spawning-capable phase. In addition, in batch spawning species, females showing postovulatory follicle complexes together with vitellogenic oocytes were also regarded as spawning capable. Individuals in this study were evaluated for assignment to the spawning-capable phase according to these criteria.

### Statistical analyses

All statistical analyses were performed using R version 4.3.1 (R Core Team 2023). Values are presented as means ± standard error of the mean. The gonadosomatic index (GSI) was calculated as GSI = (GW/BW) × 100. To assess the GSI range associated with spawning-capable status and to examine whether GSI was related to standard length (SL), SL and GSI were analyzed separately for males and females in relation to histologically determined spawning-capable status. Because neither SL nor GSI followed a normal distribution in either sex (Shapiro–Wilk test, *p* < 0.05), the relationship between SL and GSI was evaluated using Spearman’s rank correlation coefficient.

## Results

### Male testicular structure and spawning-capable phase

Of the 98 fish examined, 51 were males (SL = 182.1 ± 3.0 mm, range = 155–272 mm; GSI = 0.31 ± 0.04, range = 0.05–1.00). In males, there was no significant correlation between GSI and SL (Spearman’s ρ = 0.16, *p* = 0.27, *n* = 51; Fig. 2a). Seven of the histologically examined fish were males (SL = 203.7 ± 2.7 mm, range = 194–216 mm; GSI = 0.48 ± 0.13, range = 0.10–1.00). Across these individuals, spermatogonia, spermatocytes, spermatids, and spermatozoa were identified (Fig. 3a). The testis was lobular in organization and exhibited an unrestricted spermatogonial distribution. Spermatogonia were distributed along the length of the testicular lobules rather than being restricted to the distal terminus. Spermatozoa were present in the lumina of the lobules at spermiation. These features indicate an unrestricted spermatogonial testis type (Grier 1981; Schulz et al. 2010). Mature spermatozoa were observed in the lumina of the lobules and in the sperm ducts of all histologically examined males. Therefore, all of these individuals were inferred to be in the spawning-capable phase.

**Fig. 2.**
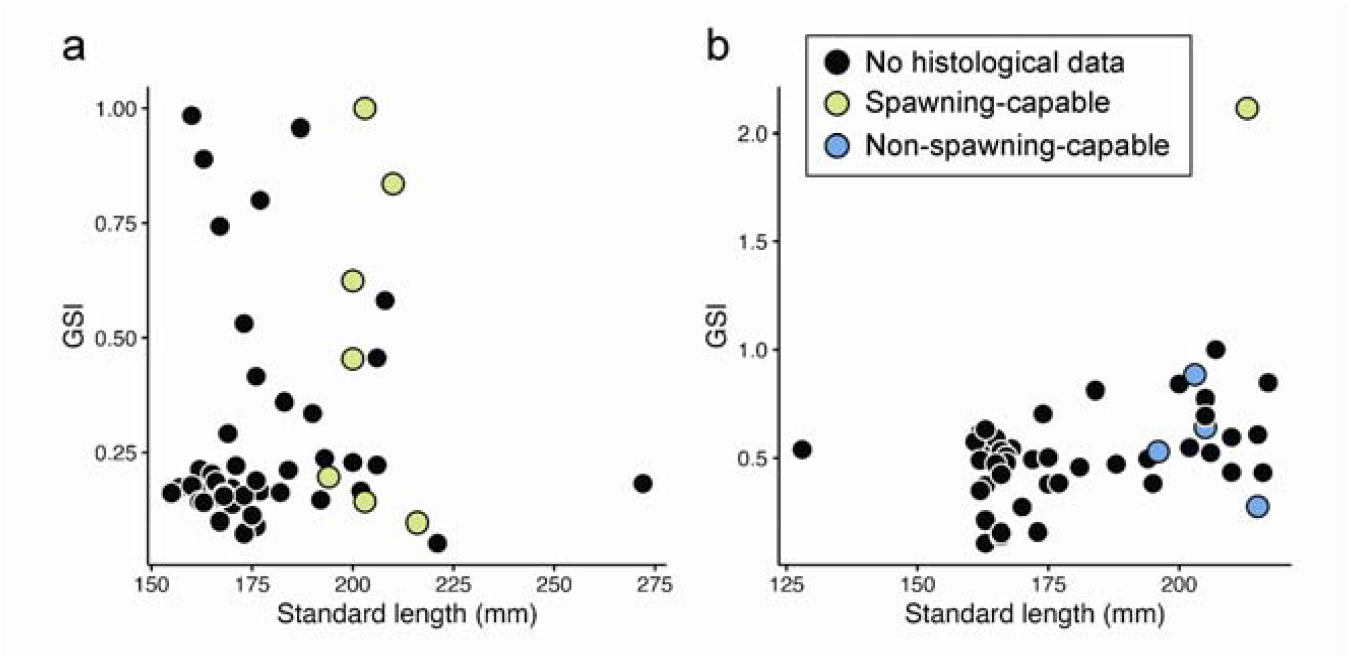
Relationship between gonadosomatic index (GSI) and standard length (SL) in male (**a**, n = 51) and female (**b**, n = 47) moonfish Mene maculata. Black points indicate specimens without histological data. Yellow and blue points indicate specimens histologically classified as spawning-capable and non-spawning-capable, respectively. No significant correlation was detected between GSI and SL in males (Spearman’s ρ = 0.16, *p* = 0.27), whereas a significant positive correlation was observed in females (Spearman’s ρ = 0.35, *p* < 0.05)

**Fig. 3.**
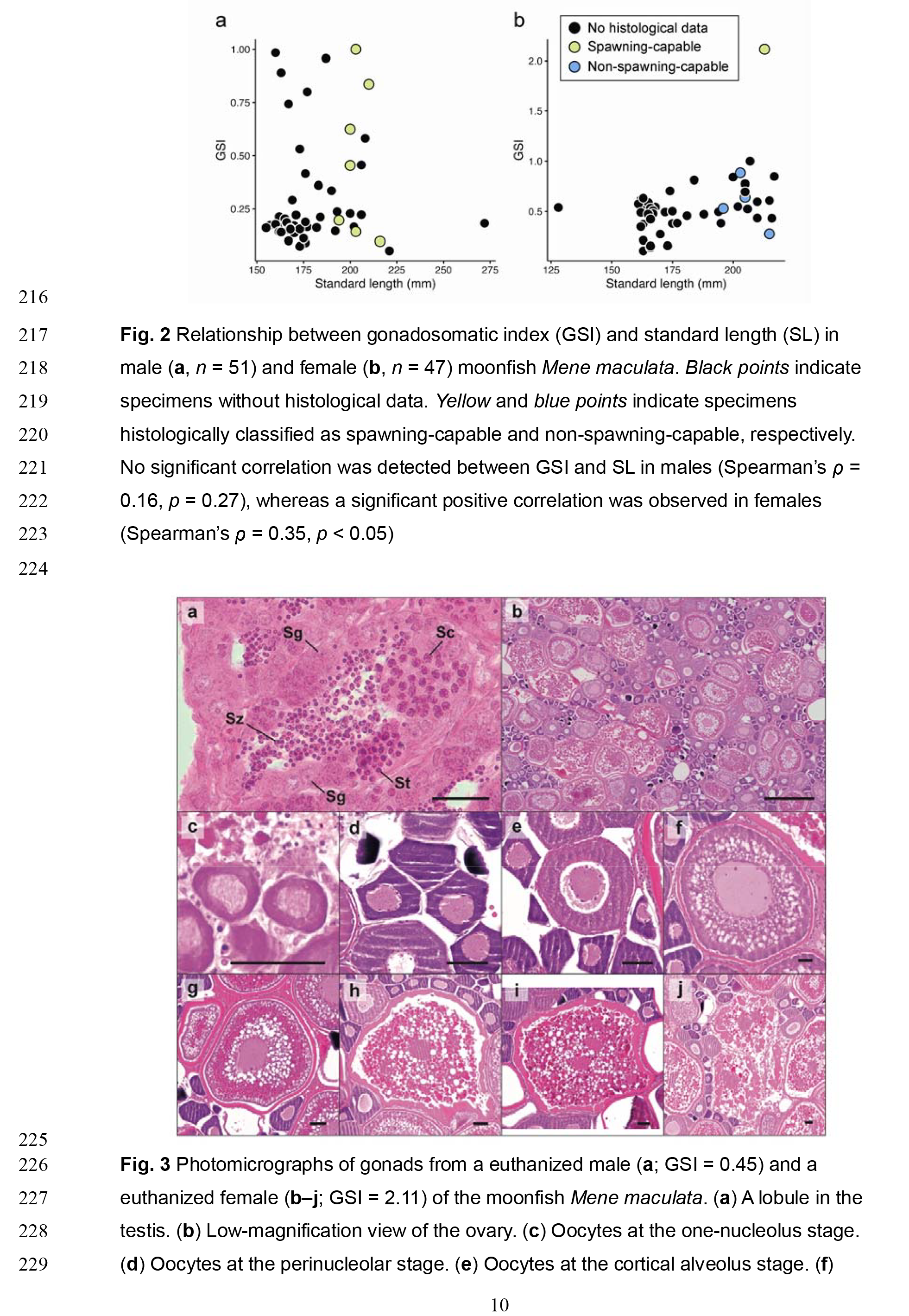
Photomicrographs of gonads from a euthanized male (**a**; GSI = 0.45) and a euthanized female (**b–j**; GSI = 2.11) of the moonfish Mene maculata. (**a**) A lobule in the testis. (**b**) Low-magnification view of the ovary. (**c**) Oocytes at the one-nucleolus stage. (**d**) Oocytes at the perinucleolar stage. (**e**) Oocytes at the cortical alveolus stage. (**f**) Oocytes at the primary vitellogenic stage. (**g**) Oocytes at the secondary vitellogenic stage. (**h**) Oocytes at the tertiary vitellogenic stage. (**i**) Oocytes at the germinal vesicle migration stage. (**j**) Atretic oocyte. Sc, spermatocyte; Sg, spermatogonia; St, spermatid; Sz, spermatozoa. Scale bars = 50 µm in **a** and **c–j**, and 500 µm in **b**

### Female ovarian structure and spawning-capable phase

Of the 98 fish examined, 47 were females (SL = 182.3 ± 3.1 mm, range = 128–217 mm; GSI = 0.54 ± 0.04, range = 0.11–2.11). In females, GSI was significantly and positively correlated with SL (Spearman’s ρ = 0.35, *p* < 0.05, *n* = 47; Fig. 2b). Five of the histologically examined fish were females (SL = 206.4 ± 3.5 mm, range = 196–215 mm; GSI = 0.89 ± 0.32, range = 0.27–2.11). Histological examination identified oocytes at the primary growth phase, including the one-nucleolus (Fig. 3c), perinucleolar (Fig. 3d), and cortical alveolus (Fig. 3e) stages; the secondary growth phase, including the primary (Fig. 3f), secondary (Fig. 3g), and tertiary (Fig. 3h) vitellogenic stages; and the oocyte maturation phase, represented by germinal vesicle migration (Fig. 3i). Atretic oocytes were also observed (Fig. 3j).

According to the criteria of Brown-Peterson et al. (2011), only the female with a GSI of 2.11 was assigned to the spawning-capable phase. In this individual, oocytes at all developmental stages from the primary growth phase to the oocyte maturation phase, as well as atretic oocytes, were observed (Fig. 3b–j). No postovulatory follicle complexes were observed in any of the females examined histologically. The simultaneous occurrence of cortical alveolus oocytes and vitellogenic oocytes at multiple developmental stages indicated asynchronous ovarian development. In the other females examined histologically, the most advanced oocytes were generally at the perinucleolar stage, although one individual contained a small number of cortical alveolus oocytes and primary vitellogenic oocytes.

## Discussion

### Testicular structure

The testis of *M. maculata* was lobular in organization and exhibited an unrestricted spermatogonial distribution. The restricted spermatogonial testis type, in which spermatogonia are confined to the distal termini of the lobules, is characteristic of Atherinomorpha, whereas the unrestricted spermatogonial testis type observed in *M. maculata* occurs widely among neoteleostean fishes (Parenti and Grier 2004; Uribe et al. 2014). Thus, although *M. maculata* is a phylogenetically distinctive species and the sole extant member of the family Menidae, no clear specialization was detected in the basic testicular organization or spermatogonial distribution examined in this study.

Although GSI spanned an approximately tenfold range among the seven males examined histologically (0.10–1.00), mature spermatozoa were present in the lumina of the lobules and sperm ducts of every individual, and all males were considered to be in the spawning-capable phase. These findings indicate that, within the range examined histologically in this study, males with relatively low GSI values can nevertheless retain mature spermatozoa and meet the histological criteria for the spawning-capable phase. Thus, in male *M. maculata*, GSI alone may have limited utility as an indicator of spawning-capable status.

In fishes, relative testis size is positively associated with the intensity of sperm competition among species, and GSI or relative testis mass is widely used as an index of relative investment in sperm production (Stockley et al. 1997; Rowley et al. 2019). In this context, the absence of a significant correlation between SL and GSI in males indicates that, within the body size range examined, larger males did not show greater relative testicular investment.

Therefore, variation in male GSI may reflect factors other than body size, such as reproductive timing, sperm production or storage, nutritional condition, and recent sperm release history. Moreover, the presence of mature spermatozoa even in low-GSI males suggests that males of this species can retain mature spermatozoa and maintain spawning-capable status without marked testicular enlargement.

### Ovarian structure

Histological examination of the ovaries obtained from the female euthanized for sampling showed asynchronous ovarian development in the only individual considered to be in the spawning-capable phase. This feature is consistent with the ovarian characteristics of batch spawning species as defined by Brown-Peterson et al. (2011), and contrasts with the total spawning pattern inferred from a unimodal oocyte diameter frequency distribution in the population off the southern coast of Java (Gani et al. 2024). Although oocyte diameter frequency distributions provide useful supplementary information on oocyte composition within the ovary, the distributional pattern alone has limited ability to determine ovarian developmental pattern and spawning pattern unambiguously. Indeed, batch spawners include not only species with asynchronous ovarian development but also species with group-synchronous ovarian development; therefore, simple unimodality or multimodality alone is likely insufficient to distinguish total spawning from batch spawning (Murua and Saborido-Rey 2003; Brown-Peterson et al. 2011). In the present study, cortical alveolus oocytes, primary, secondary, and tertiary vitellogenic oocytes, and oocytes undergoing germinal vesicle migration co-occurred within the ovary of a single individual (Fig. 3b). This histological evidence supports the possibility that *M. maculata* is a batch spawner with asynchronous ovarian development rather than a total spawner. A small number of atretic oocytes was also observed. In this individual, vitellogenic oocytes at multiple stages and oocytes undergoing germinal vesicle migration were observed simultaneously. Therefore, the few atretic oocytes were not interpreted as indicating overall regression of the ovary (Brown-Peterson et al. 2011; Corriero et al. 2021). However, this inference is based on a single female maintained in captivity. The spawning pattern of *M. maculata* therefore remains to be established through examination of additional females, including wild specimens, together with observations of spawning.

### Implications for spawning season

The spawning season of *M. maculata* has previously been inferred from seasonal changes in GSI. In the northern South China Sea, GSI of both sexes were high from August to October, and this period was inferred to be the reproductive season (Du et al. 2012). Off the southern coast of Java, on the basis of seasonal changes in GSI together with macroscopic observations of gonad maturity, the spawning season was inferred to extend from September to February, with a peak in September–October (Gani et al. 2024). These inferences rely on the interpretation that periods of relatively elevated GSI during the annual cycle correspond to advanced gonadal development and spawning-capable status. The present study complements this interpretation by histologically examining the correspondence between GSI and the spawning-capable phase in this species. The GSI of the female histologically confirmed to be in the spawning-capable phase in the present study was 2.11, which was close to the mean female GSI reported for September by Gani et al. (2024), 2.17%, during the inferred spawning peak. Therefore, the female result was broadly consistent with previous spawning season estimates based on GSI. By contrast, in males, low GSI did not necessarily indicate the absence of mature spermatozoa or loss of spawning-capable status. Therefore, estimates of the male reproductive period based on GSI require cautious interpretation. In Gani et al. (2024), the mean male GSI declined after the September–October spawning peak and reached its lowest value of 0.19% in January. This value falls within the GSI range of males considered histologically to be in the spawning-capable phase in the present study. Thus, even if the monthly mean male GSI declines after the spawning peak, it is difficult to conclude from this alone that males have left the spawning-capable phase. The present findings therefore refine previous GSI-based interpretations by showing that male GSI decline does not necessarily correspond to histological loss of spawning-capable status.

However, the present results should not be used to directly infer the spawning season of wild populations of this species. First, many of the specimens were obtained from fish maintained under controlled aquarium conditions, and the timing of their gonadal development does not necessarily correspond to that of wild populations. Second, the specimens examined here were collected from waters around Amami-Oshima, near the northern margin of the species’ distribution, and therefore may not represent the general seasonal reproductive cycle of this species in the central part of its range. Previous studies investigated GSI variation in populations from the South China Sea and off the southern coast of Java, and regional or latitudinal differences in spawning timing may exist. At the same time, the present study shows that interpretation of GSI and spawning-capable status in this species should take into account sex-specific histological relationships, and provides histological evidence that complements the GSI-based spawning season estimates of Du et al. (2012) and Gani et al. (2024). Future studies integrating year-round histological surveys of wild populations with observations of spawning behavior would provide a more comprehensive understanding of the reproductive ecology of this species.

## Acknowledgments

We are grateful to T. Ueda, H. Imura, and I. Takehara for collecting live specimens. We also thank A. Komoda for valuable advice and helpful discussions, and the aquarium staff for their assistance throughout the study. We thank K. Ohara and M. Ando for assistance with the investigation. This study was funded by the Marine Science Museum Fukushima, Mikimoto Fund for Marine Ecology, and Japan Society for the Promotion of Science (JSPS) KAKENHI Grant Numbers 24932659, 23721203, and 24940220.

## Declarations

### Ethics approval

At Kitasato University, there are currently no institutional ethical regulations specifically for research on fish. Therefore, all procedures were conducted in accordance with the Guidelines for the ethical treatment of nonhuman animals in behavioural research and teaching of the Association for the Study of Animal Behaviour and Animal Behavior Society (ASAB Ethical Committee/ABS Animal Care Committee 2023), as well as the Guidelines for the use of fishes in research published by the Ichthyological Society of Japan in 2003 (https://www.fish-isj.jp/message/guideline/2003_1011/).

### Data availability

The datasets are available from the corresponding author on reasonable request.

### Declaration of generative AI use

During manuscript preparation, the authors used ChatGPT (OpenAI) to assist with

English-language drafting and revision. The authors reviewed and revised all AI-assisted text and take full responsibility for the accuracy and integrity of the final manuscript.

